# The mating type transcription factor MAT1-1-1 from the fungal human pathogen *Aspergillus fumigatus*: synthesis, purification, and crystallization of the DNA binding domain

**DOI:** 10.1101/2021.12.13.472399

**Authors:** Barbara Ramšak, Ulrich Kück, Eckhard Hofmann

## Abstract

Mating type (*MAT*) loci are the most important and significant regulators of sexual reproduction and development in ascomycetous fungi. Usually, they encode two transcription factors (TFs), named MAT1-1-1 or MAT1-2-1. Mating-type strains carry only one of the two TF genes, which control expression of pheromone and pheromone receptor genes, involved in the cell-cell recognition process. The present work presents the crystallization for the alpha1 (α1) domain of MAT1-1-1 from the human pathogenic fungus *Aspergillus fumigatus* (AfMAT1-1-1). Crystals were obtained for the complex between a polypeptide containing the α1 domain and DNA carrying the AfMAT1-1-1 recognition sequence. A streak seeding technique was applied to improve native crystal quality, resulting in diffraction data to 3.2 Å resolution. Further, highly redundant data sets were collected from the crystals of selenomethionine-substituted AfMAT1-1-1 with a maximum resolution of 3.2 Å. This is the first report of structural studies on the α1 domain MAT regulator involved in the mating of ascomycetes.

**Synopsis:** An optimized purification and crystallization protocol together with initial X-ray datasets are described for this mating type transcription factor from human pathogenic fungus *Aspergillus fumigatus*.

## Introduction

*Aspergillus fumigatus* is one of the most prevalent pathogenic fungi in humans, causing pulmonary aspergillosis in immunocompromised individuals with a reported mortality rate of 30-95% (Brown *et al*., 2012). Rising trends in the incidence of aspergillosis are related to an increased use of immunosuppressive drugs, and to an emergence of *A. fumigatus* strains resistant to azole antifungal-drugs (van der Linden *et al*., 2015). Together with the recently reported cases of COVID-19-associated pulmonary aspergillosis caused by azole-resistant aspergillus, they pose a serious global health concern (Arastehfar *et al*., 2020).

The adaptive potential of *A. fumigatus* to colonize changing or new habitats is linked to the sexual reproduction that was recently described (O’Gorman *et al*., 2009; Zhang *et al*., 2017). Two distinct mating types determined by two non-allelic mating-type loci, named *MAT1-1* and *MAT1-2*, are a prerequisite for entry into mating process (Paoletti *et al*., 2005; Szewczyk & Krappmann, 2010). These loci encode either the MAT1-1-1 or the MAT1-2-1 transcription factor (TF) that were found to regulate expression of numerous genes involved in sexual and cellular development in diverse ascomycetes (Becker *et al*., 2015; Kim *et al*., 2015; Wada *et al*., 2012; Yu *et al*., 2018).

MAT1-1-1 contains a conserved ∼40 amino acid alpha1 (α1) domain consisting of four alpha helices (Martin *et al*., 2010). This domain is also present in the well-characterized Mat1α (MAT1-1-1) mating-type TF from the yeast *Saccharomyces cerevisiae*, which was used previously as a model to get a mechanistic understanding of eukaryotic signal transduction pathways (Bardwell, 2005; Engelberg *et al*., 2014). Recently, we characterized the DNA binding motif of MAT1-1-1 derived from promoter regions of several developmentally regulated genes, including the one for the pheromone receptor PreA (Ramšak *et al*., 2021). Several attempts have been undertaken to crystallize MAT1-1-1-like proteins (Jackson *et al*., 2013; Dyer *et al*., 2016), however no successful report has been published anywhere. Here we report for the first time the purification and crystallization protocol that results in the formation of crystals of a fungal α1 domain in complex with its target preA-2 DNA.

### Experimental procedures

#### Macromolecule production

Plasmid vector pGEX-AfMAT1-1-1_F5 for inducible expression of recombinant AfMAT1-1-1 protein (residues 78-235) has previously been reported (Ramšak *et al*., 2021). The vector contains a GST-affinity tag fused to the region encoding N-terminus of the AfMAT1-1-1_78-235_. The sequence encoding the tobacco etch virus (TEV) protease cleavage site separates the GST-tag and the AfMAT1-1-1_78-235_ coding sequence, leaving three additional residues (GMA) at the N-terminus. The AfMAT1-1-1_78-235_ synthesis and purification was performed as described in detail in Ramšak *et al*. (2021). Macromolecule-production information is summarized in **Table 1**, and the buffer compositions used for purification are listed in **Table S1**.

**Table 1.** Information on the AfMAT1-1-1_78-235_.

|  |  |
| --- | --- |
| Source organism | *Aspergillus fumigatus* (strain Af293) |
| Expression vector | pGEX-AfMAT1-1-1_F5 |
| Host | *E. coli* strain BL21 Gold(DE3) pLysS |
| Complete amino-acid sequence of the construct produced† | <u>GAM</u>AKRTQEGKKRPLNSFIAFRSFYSVIFPDLTQKAKSGILRFLWQND<br>PF<br>KAKWAILAKAYSII RDDHESEVSLDQFLEITAKFIGLFEP<br>A<br>RYLDAMGWQLNFDDQQQYTM AKVKITTIPEADVSTNYSV<br>GDIVKHCYDTGYVSEKPGKHTGSNGNNTSTM |

† The underlined sequence originates from the linker between GST-tag and the AfMAT1-1-1<sub>78-235</sub> polypeptide.

#### Crystallization of AfMAT1-1-1_78-235_ in complex with preA-2 DNA

For crystallization trials, the complementary 29 bp oligonucleotides preA-2_f and preA-2_r (Ramšak *et al*., 2021) were purchased from Sigma-Aldrich in HPLC purity grade. The preA-2 DNA duplex consisted of the -CTATTGAG-consensus motif and 5’-AT overhang (**Fig. 1**) and was previously shown to be bound by the truncated AfMAT1-1-1_78-235_ (Ramšak *et al*., 2021). Prior to crystallization the complementary strands were annealed in equimolar ratios in 10 m*M* Tris pH 7.5, 50 m*M* NaCl, 1 m*M* EDTA, to a final concentration of 2.7 m*M*. AfMAT1-1-1_78-235_ and the annealed DNA duplex were mixed in 2:1 molar ratio. Next, the protein-DNA complex was incubated for 30 min on ice. For crystallization experiments, the high salt content of the protein buffer was lowered by dialysis against 50 m*M* HEPES pH 8.0, 150 m*M* NaCl, 2 m*M* DTT to optimize binding of the protein with oligonucleotides.

**Figure 1.**
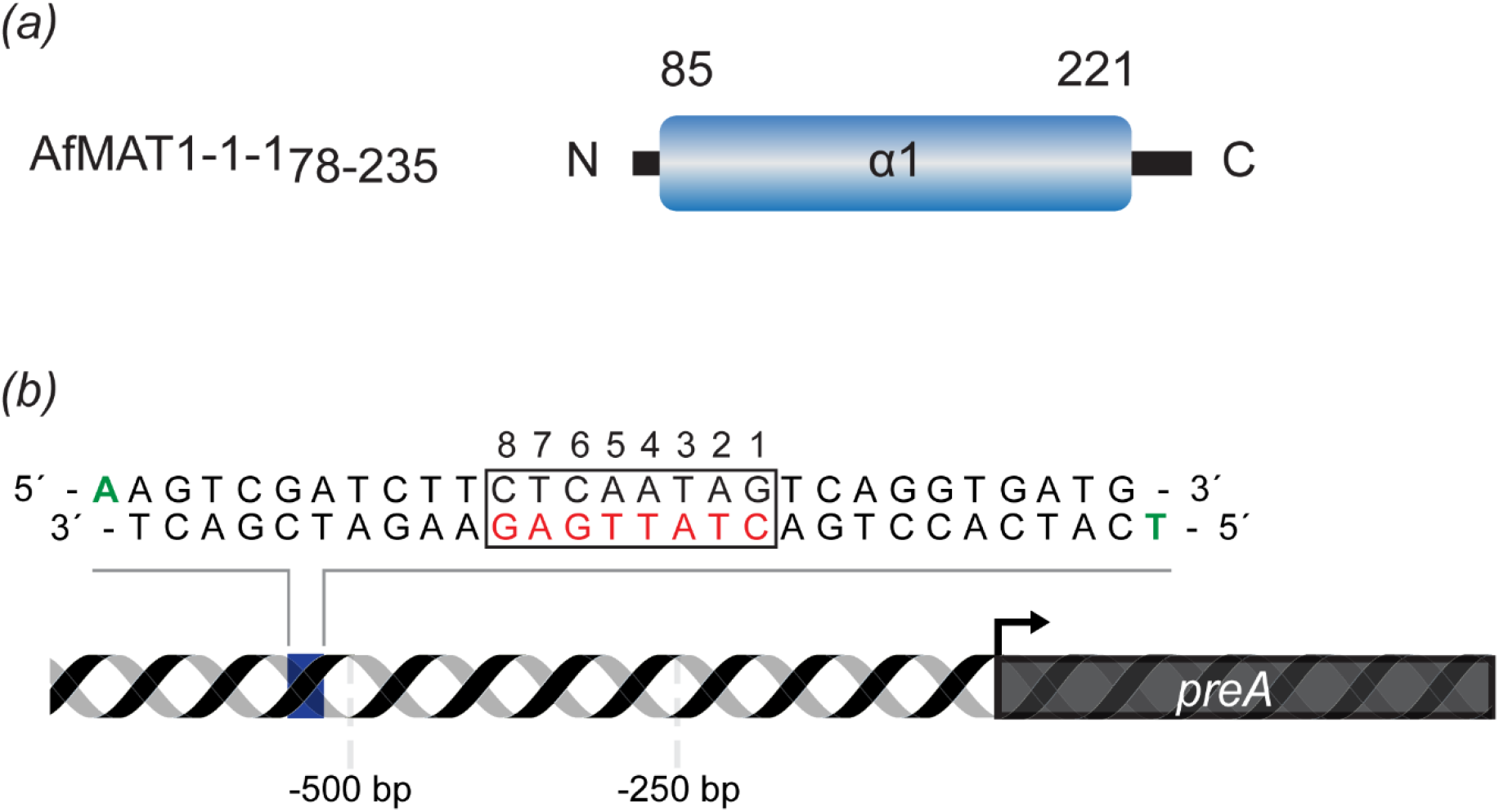
Schematic representation of the AfMAT1-1-1_78-235_ and DNA (preA-2) used for co-crystallization. (*a*) The AfMAT1-1-1 construct consisting of α1 domain, delimited by residues numbers (*b*) Sequence of the 29-mer DNA (preA-2) was derived from the *preA* promoter region and carries the DNA binding site (Ramšak *et al*., 2021). The binding sequence for AfMAT1-1-1 is boxed. Nucleotides colored in green denote A and T overhangs.

Preliminary crystallization screening was carried out by sitting drop vapour diffusion using a Crystal Phoenix (Art Robbins Instruments) robotic system, mixing 100 nl protein with 100 nl reservoir solution. In total 240 unique buffer conditions were tested, which were derived from commercially available screens Nucleix (NeXtal), protein-nucleic acid complex crystal screen from Kerafast (Pryor *et al*., 2012), and the PEGII Suite (Qiagen). The PEGII Suite (Qiagen) was included since it was previously reported that the protein-DNA complexes preferentially crystalize in a wide-range of polyethylene glycol (PEG) molecular weights (Dock-Bregeon *et al*., 1999; Pryor *et al*., 2012). Crystals in the initial hits appeared after 3 days and continued to grow for 7 days (**Fig. 2**). The best hits were observed in two conditions: a) 0.1 *M* Tris-HCl pH 8.0, 30% (w/v) PEG 4000, 2.0 *M* LiSO4 (PEGII Suite) and b) 0.1 *M* HEPES pH 7.5, 25% (w/v) PEG 2000, 0.1 *M* CaCl_2;_ 0.1 *M* NaCl (Pryor *et al*., 2012). Conditions were subsequently refined in hanging-drop settings and are summarized in **Table 2**. Presence of both the protein and preA-2 DNA in the crystals was confirmed (**Fig. 3**) by washing 10-15 crystals in the three separate drops of reservoir solution and dissolving them as described in Pryor *et al*. (2012).

**Table 2.** Crystallization.

| Construct | AfMAT1-1-1<sub>78-235</sub> | SeMet-AfMAT1-1-1<sub>78-235</sub> |
| --- | --- | --- |
| Method | Hanging-drop vapor diffusion | Hanging-drop vapor diffusion |
| Plate type | VDX plates (Hampton Research) | VDX plates (Hampton Research) |
| Temperature (K) | 291 | 291 |
| Protein | 9-10 mg ml<sup>-1</sup><br>2:1 (protein:dsDNA) | 9-10 mg ml<sup>-1</sup><br>2:1 (protein:dsDNA) |
| Buffer composition of protein solution | 50 mM HEPES pH 8.0, 150 mM NaCl, 2 mM DTT | 50 mM HEPES pH 8.0, 150 mM NaCl, 2 mM DTT |
| Composition of refined reservoir solution | 0.1 M Tris-HCl pH 8.0, 26% (w/v) PEG 4000, 1.2 M LiSO<sub>4</sub> | 0.1 M HEPES pH 7.5, 22.5% (w/v) PEG 2000, 0.15 M CaCl<sub>2</sub>, 0.1 M NaCl |
| Volume and ratio of drop | 2 µl:2 µl (protein:reservoir)<br>protein:reservoir solution | 2 µl:2 µl (protein:reservoir)<br>protein:reservoir solution |
| Volume of reservoir (µl) | 700 | 700 |

**Figure 2.**
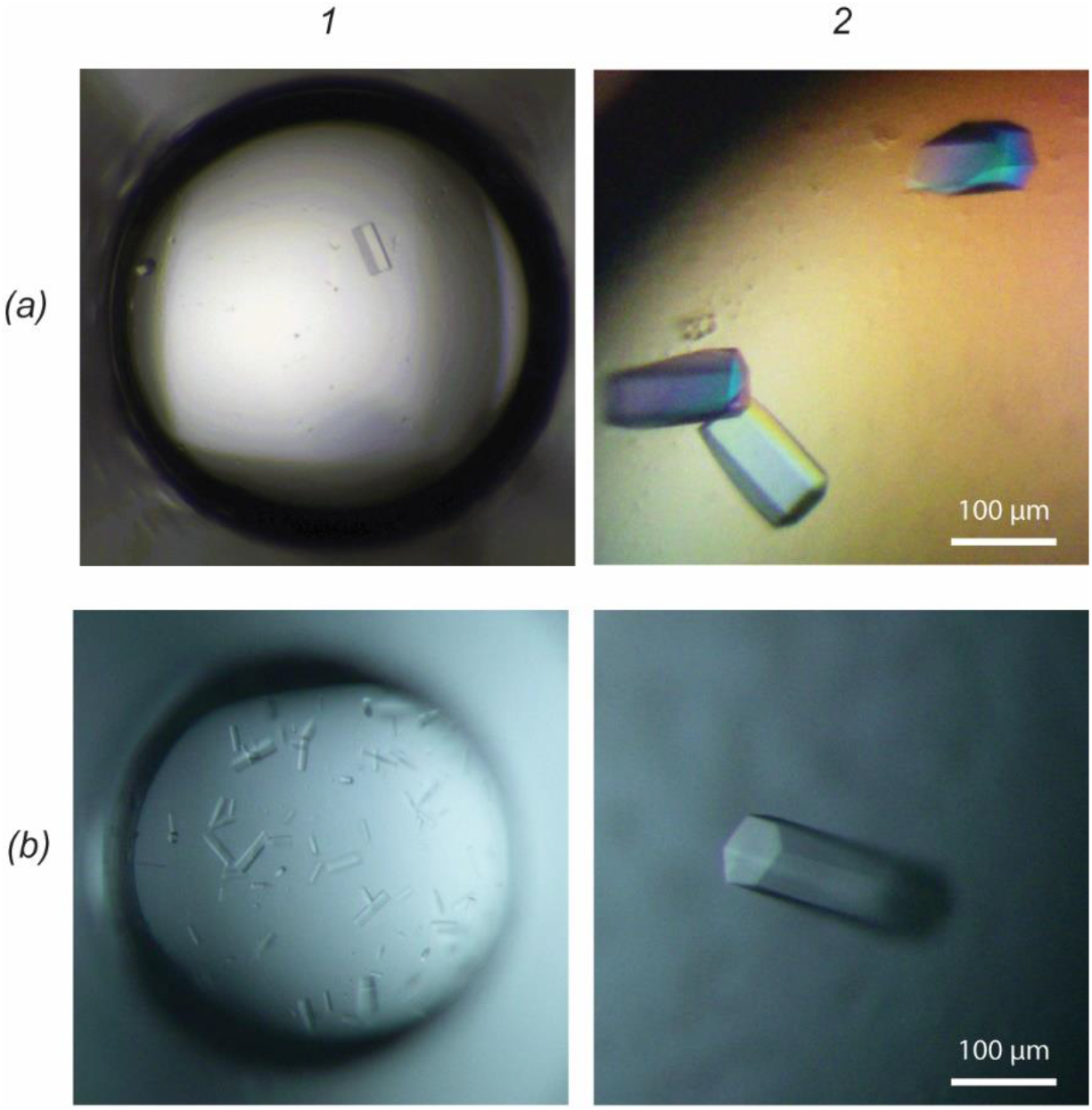
Hexagonal crystals of the AfMAT1-1-1_78-235_/preA-2 complex. *(a1)* Native crystals grown in sitting-drop, and in hanging-drop *(a2)* obtained after streak seeding in a refined buffer condition. SeMet crystals grown in sitting-drop *(b1)* and in hanging-drop *(b2)* in a refined buffer condition.

**Figure 3.**
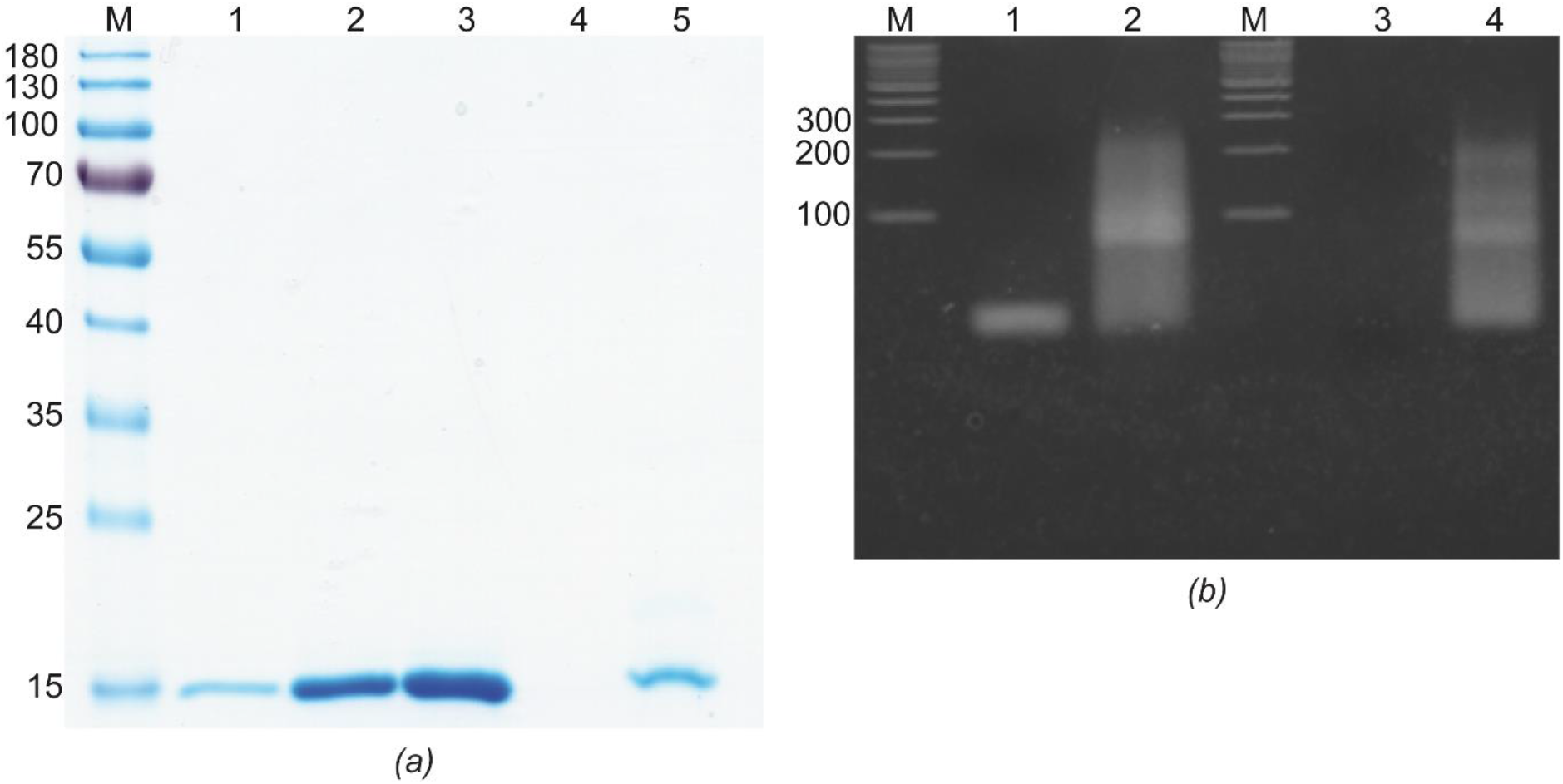
Characterization of the AfMAT1-1-1_78-235_/preA-2 complex crystal by electrophoresis. (*a*) SDS-PAGE analysis of the purified AfMAT1-1-1_78-235_ (18 kDa) used in co-crystallization trials, and dissolved native crystals. Lane *M*, molecular-weight marker (in kDa); lane *1-3*, 1, 5 and 10 μg of purified AfMAT1-1-1_78-235_, respectively; Lane *4*, drop used for crystal wash; Lane *5*, several dissolved crystals. 12% SDS-PAGE was stained with Coomassie Blue Stain. (*b*) 4% agarose gel electrophoresis analysis. Lane *M*, DNA weight marker (in bp); Lane *1*, annealed preA-2 DNA; lane *2* AfMAT1-1-1_78- 235_/preA-2 complex before crystallization; lane 3, as a control washing buffer for crystals before solving was loaded; lane *4*, several dissolved crystals. Gel is stained with ethidium bromide.

#### X-ray data collection and processing

Crystals were flash-cooled in liquid nitrogen after being cryoprotected with reservoir solution containing 15% PEG 400. X-ray diffraction data were collected at the European Synchrotron Radiation Facility (ESRF, Grenoble, France) on the beamlines ID30B and ID32-2 for native and SeMet-protein/DNA complex crystals, respectively (**Fig. 4**). Multiple data sets at different crystal positions and with different κ angles were collected for the SeMet-crystal complex. Processing and scaling of data sets was carried out with the *XDS* (Kabsch, 2010) software suite. Data collection and initial processing statistics are summarized in **Table 3**.

**Table 3.** Crystal data collection and processing. Values in parentheses are for the highest resolution shell.

|  | Native | SeMet derivative<sup>†</sup> |
| --- | --- | --- |
| Diffraction Source | ESRF ID30B | ESRF ID23-2 |
| Wavelength (Å) | 0.975490 | 0.873130 |
| Temperature (K) | 100 | 100 |
| Detector | PILATUS3 6M | PILATUS3 2M |
| Crystal-to-detector distance (mm) | 578.5 | 411.1, 815.0, 323.0<sup>††</sup> |
| Rotation range per image (°) | 0.1 | 0.1 |
| Total rotation range (°) | 360 | 14*360 |
| Exposure time per image (s) | 0.02 | 0.02 |
| Flux (photon/s) | 1.8*10<sup>12</sup> | 1.7*10<sup>11</sup> |
| Space group | *P*6<sub>2</sub>22 or *P*6<sub>4</sub>22 | *P*3<sub>1</sub>21 |
| a, b, c (Å) | 91.28, 91.28, 271.55 | 89.49, 89.49, 270.86 |
| α, β, γ (°) | 90, 90, 120 | 90, 90, 120 |
| Resolution range (Å) | 45.64-3.20 (3.28-3.20) | 44.74-3.20 (3.28-3.20) |
| Total No. of reflections | 428951 (32598) | 5066659 (369303) |
| No. of unique reflections | 11822 (852) | 39977 (2959) |
| Completeness (%) | 99.9 (100) | 99.9 (100) |
| Multiplicity | 36.2 (38.2) | 126.7 (124.8) |
| ⟨I/σ(I)⟩ | 19.72 (1.06) | 21.45 (0.81) |
| R<sub>meas</sub> (%) | 12.6 (418.2) | 24 (504.8) |
| CC1/2 | 99.9 (55.6) | 100.0 (75.9) |
| Overall B factor from Wilson plot (Å<sup>2</sup>) | 129.8 | 120.5 |

<sup>†</sup>14 sweeps, 13 merged; see methods.<sup>††</sup>Most sweeps were collected with a detector distance of 411.1 mm.

**Figure 4.**
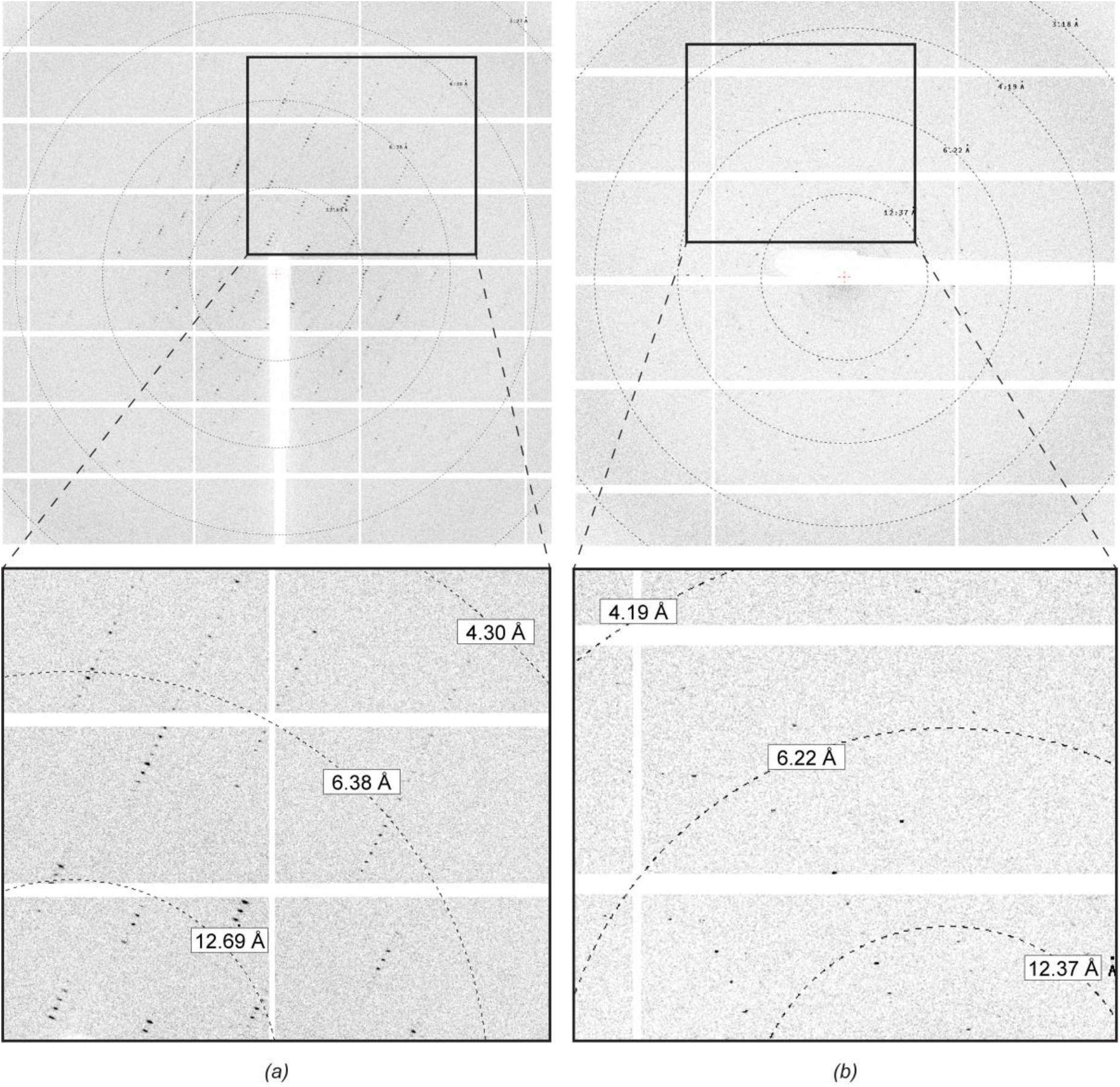
Representative X-ray diffraction patterns. (a) 0.1° oscillation image from the native crystal, collected on beamline ID30B at the ESRF. (b) 0.1° oscillation image from the SeMet-substituted crystal, collected at beamline ID23-2 at the ESRF.

## Results and discussion

AfMAT1-1-1_78-235_ (Mw = 18 000 Da) was purified to >95% purity as determined by SDS–PAGE analysis with typical yields of 2.5-3.5 mg pure protein per litre of bacterial expression culture (**Fig. 3a**). Addition of charged amino acids as described above proved to be essential in protein preparation. A great effort has been made to produce the diffraction-quality crystals of AfMAT1-1-1_78-235_/preA-2 complex (**Fig. 3**). The optimal protein:DNA ratio significantly affected formation of the crystal complexes, as no crystals were obtained when protein and DNA were mixed in ratio 1:1.2 or when AfMAT1-1-1_78-235_ alone was used for crystallization screenings. Diffraction data collected for native crystals was limited to ∼5-8 Å resolution, with poor reproducibility. Therefore, the streak-seeding technique according to Stura and Wilson (1991) was applied to obtain better ordered crystals. Microseeds obtained from crushed crystals grown in conditions with 30% PEG 4000 were transferred with a cat whisker to freshly prepared drops of non-nucleated protein with 26% PEG 4000 (**Fig. 2b**). Crystals prepared by this approach were not only more stable, but also diffracted up to 3.2 Å resolution. It is worth mentioning that the optimization of the cryobuffers by using diverse cryoprotectants such as glycerol and sucrose did not improve data quality further. A complete data set was collected for a PEG 400 cryoprotected native crystal from seeding set-up, belonging to space group *P*6_2_22 or *P*6_4_22 with unit-cell parameters a=b=91.28, c= 271.55. Analysis of the Matthews coefficient (Weichenberger & Rupp, 2014) suggests 2 copies of TF with dsDNA in the asymmetric unit (Vm=2.6 Å^3^/Da, Vs=57%).

The amino acid sequence for the α1 domain shows only weak homology (28% identity) to previously published sequences for high-mobility group box (HMG-box) domains, which are regularly found in MAT1-2-1 TFs from filamentous ascomycetes (Martin *et al*., 2010). Accordingly, initial attempts to solve the structure by molecular replacement failed. We therefore screened a SeMet-AfMAT1-1-1_78- 235_/preA-2 complex to obtain experimental phase information. Interestingly, the initial SeMet crystals were obtained only under condition b) mentioned above. The refinement of this condition resulted in one single crystal (**Fig. 2b**) with space group *P*3_1_21 and unit-cell parameters a=b=89.49, c=270.86, for which multiple data sets were collected to 3.2 Å resolution. Streak seeding was performed for SeMet crystals as well, however this did not improve diffraction further. Analysis of the Matthews coefficient (Weichenberger & Rupp, 2014) suggests 4 copies of TF with dsDNA in the asymmetric unit (Vm=2.75 Å^3^/Da, Vs=59%). Anomalous signal was present to 3.7 Å in the high-redundancy dataset. Substructure determination and phasing using HKL2MAP and the SHELX-suite was attempted (Pape & Schneider, 2004; Sheldrick, 2008). While SHELXD clearly identified 8 sites with occupancy above 50%, the resulting phasing with SHELXE did not yield phases of sufficient quality for model building, most likely due to the moderate resolution of the data. Attempts utilizing other phasing pipelines were unsuccessful so far.

Currently, we are working on improving resolution and phasing to obtain the structure of this TF/DNA complex. The strategy described here could be applied for other TFs from filamentous fungi for which so far only very limited crystallization reports are available.

## Supporting information

RamsakEtal-Suppl-Tab1

## Acknowledgements

We thank the European Synchrotron Radiation Facility (ESRF) for beamtime access and the support during X-ray data collection. We are grateful to Petros Sarantopoulos for assistance with the crystallization robot and Sebastian Österlin for his help with initial crystallization preparation and crystal mounting. We thank Roman Sakson and Prof. Dr. Albert Sickmann (ISAS, Dortmund, Germany) for acquiring MS data, and Prof. Dr. Sven Krappmann (Erlangen; Germany) for fruitful collaboration. This work was supported by the German Research Foundation (DFG) (Bonn Bad-Godesberg, Germany) (KU 517/ 15-1). Part of this research was funded by the DFG Research Training Group GRK 2341 “Microbial Substrate Conversion (MiCon)”.

## Notes

**Funding information** German Research Foundation (DFG) (Bonn Bad-Godesberg, Germany) (contract No. KU 517/15-1); Research Training Group [Microbial Substrate Conversion (MiCon)] (grant No. GRK 2341).

### Competing Interest Statement

The authors have declared no competing interest.

