## Supplementary material for "The mating type transcription factor MAT1-1-1 from the fungal human pathogen *Aspergillus fumigatus*: synthesis, purification, and crystallization of the DNA binding domain": RamsakEtal-Suppl-Tab1

**Supporting information**

Table S1 Compositions of buffer used in purification of AfMAT1-1-1_78-235_.

| **Purification step** | **Buffer** | **Buffer composition** |
| --- | --- | --- |
| I | Lysis buffer | 500 m*M* NaCl, 27 m*M* KCl, 10 m*M* Na_2_HPO_4_ pH 7.3, 18 m*M* KH_2_PO_4_, 5 m*M* DTT, 0.1 % Protease-Inhibitor Cocktail IV, 0.1% 100 m*M* PMSF, DNase |
|  | Wash buffer1 | 500 mM NaCl, 27 m*M* KCl, 10 m*M* Na_2_HPO_4_ pH 7.3, 18 m*M* KH_2_PO_4_, 5 m*M* DTT |
|  | Elution buffer1 | 500 mM NaCl, 50 m*M* Tris pH 8.0, 20 m*M* reduced glutathione, 5 m*M* DTT |
| Cleavage | Cleavage buffer | 500 mM NaCl, 50 m*M* HEPES pH 7.5, 1 m*M* EDTA, 2 m*M* DTT, 50 m*M* L-Arg + L-Glu |
| II | Wash buffer2 | 500 mM NaCl, 50 m*M* HEPES pH 8.0, 1 m*M* EDTA, 30 m*M* reduced glutathione, 2 m*M* DTT, 50 m*M* L-Arg + L-Glu |
|  | Elution buffer2 | 500 m*M* NaCl, 50 m*M* HEPES pH 7.5, 1 m*M* EDTA, 2 m*M* DTT |
| III | Size-exclusion buffer | 500 m*M* NaCl, 50 m*M* HEPES pH 7.5, 1 m*M* EDTA, 2 m*M* DTT, 50 m*M* L-Arg + L-Glu |
